# Viral E Protein Neutralizes BET Protein-Mediated Post-Entry Antagonism of SARS-CoV-2

**DOI:** 10.1101/2021.11.14.468537

**Authors:** Irene P. Chen, James E. Longbotham, Sarah McMahon, Rahul K. Suryawanshi, Jared Carlson-Stevermer, Meghna Gupta, Meng Yao Zhang, Frank W. Soveg, Jennifer M. Hayashi, Taha Y. Taha, Victor L. Lam, Yang Li, Zanlin Yu, Erron W. Titus, Amy Diallo, Jennifer Oki, Kevin Holden, QCRG Structural Biology Consortium, Nevan Krogan, Danica Galonić Fujimori, Melanie Ott

## Abstract

Inhibitors of Bromodomain and Extra-terminal domain (BET) proteins are possible anti-SARS-CoV-2 prophylactics as they downregulate angiotensin-converting enzyme 2 (ACE2). Here, we show that BET proteins should not be inactivated therapeutically as they are critical antiviral factors at the post-entry level. Knockouts of BRD3 or BRD4 in cells overexpressing ACE2 exacerbate SARS-CoV-2 infection; the same is observed when cells with endogenous ACE2 expression are treated with BET inhibitors during infection, and not before. Viral replication and mortality are also enhanced in BET inhibitor-treated mice overexpressing ACE2. BET inactivation suppresses interferon production induced by SARS-CoV-2, a process phenocopied by the envelope (E) protein previously identified as a possible “histone mimetic.” E protein, in an acetylated form, directly binds the second bromodomain of BRD4. Our data support a model where SARS-CoV-2 E protein evolved to antagonize interferon responses via BET protein inhibition; this neutralization should not be further enhanced with BET inhibitor treatment.

## INTRODUCTION

The bromodomain and extraterminal domain (BET) family of proteins consists of BRD2, BRD3, BRD4, and BRDT, the latter only found in testis. BET proteins characteristically harbor two highly conserved N-terminal bromodomains (BDs [BD1 and BD2]) and an extraterminal (ET) domain. BDs function as *bone fide* reader domains of acetylated lysines in histone and non-histone proteins and are the molecular targets of small-molecule BET inhibitors such as JQ1, while the ET domain has less defined protein binding properties (Dhalluin et al., 1999; Filippakopoulos et al., 2012; Rahman et al., 2011). Through their interaction with histones and cellular transcriptional machinery, BET proteins play an instrumental role in many cellular functions, including cell proliferation, chromatin remodeling, and gene expression (Taniguchi, 2016). BRD4 is the best studied BET protein and exists in different splice isoforms: a so-called long isoform (amino acids [aa] 1-1362, BRD4L), a short isoform (aa 1-720, BRD4S), and an intermediate third isoform reported only in osteosarcoma cells (Floyd et al., 2013). BRD4L has an extended C-terminus that recruits the positive transcription elongation factor (PTEF-b) termed the PTEF-b binding domain (PID) (Bisgrove et al., 2007). Furthermore, BRD2, BRD3, and BRD4 interact with viral proteins of herpesviruses, flaviviruses, and papillomaviruses and regulate the integration and latent viral infection of retroviruses (Conrad et al., 2017; De Rijck et al., 2013; Mourao et al., 2020; Platt et al., 1999; Sharma et al., 2013; Wu et al., 2006; You et al., 2006).

The recently emerged SARS-CoV-2 virus is the causative agent of the on-going coronavirus disease 2019 (COVID-19) pandemic (Zhu et al., 2020). COVID-19 patients are characterized by impaired type I interferon (IFN-I) responses paired with an overproduction of proinflammatory cytokines (Blanco-Melo et al., 2020; Hadjadj et al., 2020; Jose and Manuel, 2020). A potent coactivator of proinflammatory and antiviral genes is BRD4. In the lung, BRD4 coactivates interferon-stimulated genes (ISGs) during viral infection by recruiting P-TEFb and activating proinflammatory responses associated with lung fibrosis, chronic obstructive pulmonary disease, and asthma (Hargreaves et al., 2009; Stratton et al., 2017). Treatment with BET inhibitors like JQ1 attenuates transcriptional activation of the antiviral response in the context of influenza A infection (Wienerroither et al., 2014). In addition, we previously identified BRD4 and BRD2 as high-confidence interactors of the SARS-CoV-2 E protein and described a histone-like motif similar to histone H3 within E protein that may interfere with the canonical histone:BET protein interactions (Gordon et al., 2020).

Notably, BRD2 functions as a transcriptional regulator of the viral entry receptor angiotensin-converting enzyme 2 (ACE2), where knockout of BRD2 or prophylactic application of BET inhibitors reduces ACE2 expression and *de novo* viral infection (Mills et al., 2021; Qiao et al., 2020; Tian et al., 2021). To reconcile this proviral function of BRD2 with known antiviral effects of BRD4, we tested the function of all relevant BET proteins during SARS-CoV-2 infection. Inhibition of BET proteins after viral entry or knockout of BRD3 or BRD4 in cells overexpressing ACE2 significantly enhanced viral replication, indicating a post-entry antiviral function of BET proteins on SARS-CoV-2. This enhancement in viral replication was also observed in infected, BET inhibitor-treated K18-hACE2 mice. The viral E protein in an acetylated form effectively thwarts this antiviral function as it binds to the second bromodomain of BRD4, underscoring the relevance of BET proteins as positive regulators of antiviral gene expression and cautioning against the use of BET inhibitors in an ongoing SARS-CoV-2 infection.

## RESULTS

### BRD3 and BRD4 Proteins Antagonize SARS-CoV-2 Infection

To test post-entry functions of all relevant BET proteins, we knocked them out in A549 cells overexpressing ACE2 (A549-ACE2). We generated polyclonal knockout (KO) cells of BRD2, BRD3, or BRD4 via nucleofection of CRISPR-Cas9 ribonucleoprotein (RNP) complexes incorporating multiple guide RNAs against one target. As controls, cells were nucleofected with the RNP complex without guide RNAs (RNP only). The knockout of individual BET proteins was validated by western blotting before infection with SARS-CoV-2 (Isolate USA/WA-1/2020) at multiplicities of infection (MOI) of 0.01 and 0.1 (Figure 1A). Loss of BRD3 and BRD4 significantly enhanced cell-associated viral RNA titers (9-fold and 17-fold, respectively) and infectious particle production in plaque assays (1.4 log fold increase for BRD4) at MOI of 0.1, while only BRD4 KO significantly enhanced viral replication in cells infected at MOI of 0.01 (Figure 1B and 1C). BRD2 KO did not significantly enhance viral infection at any MOI supporting a model where individual BET proteins play distinct roles in SARS-CoV-2 infection.

**Fig 1.**
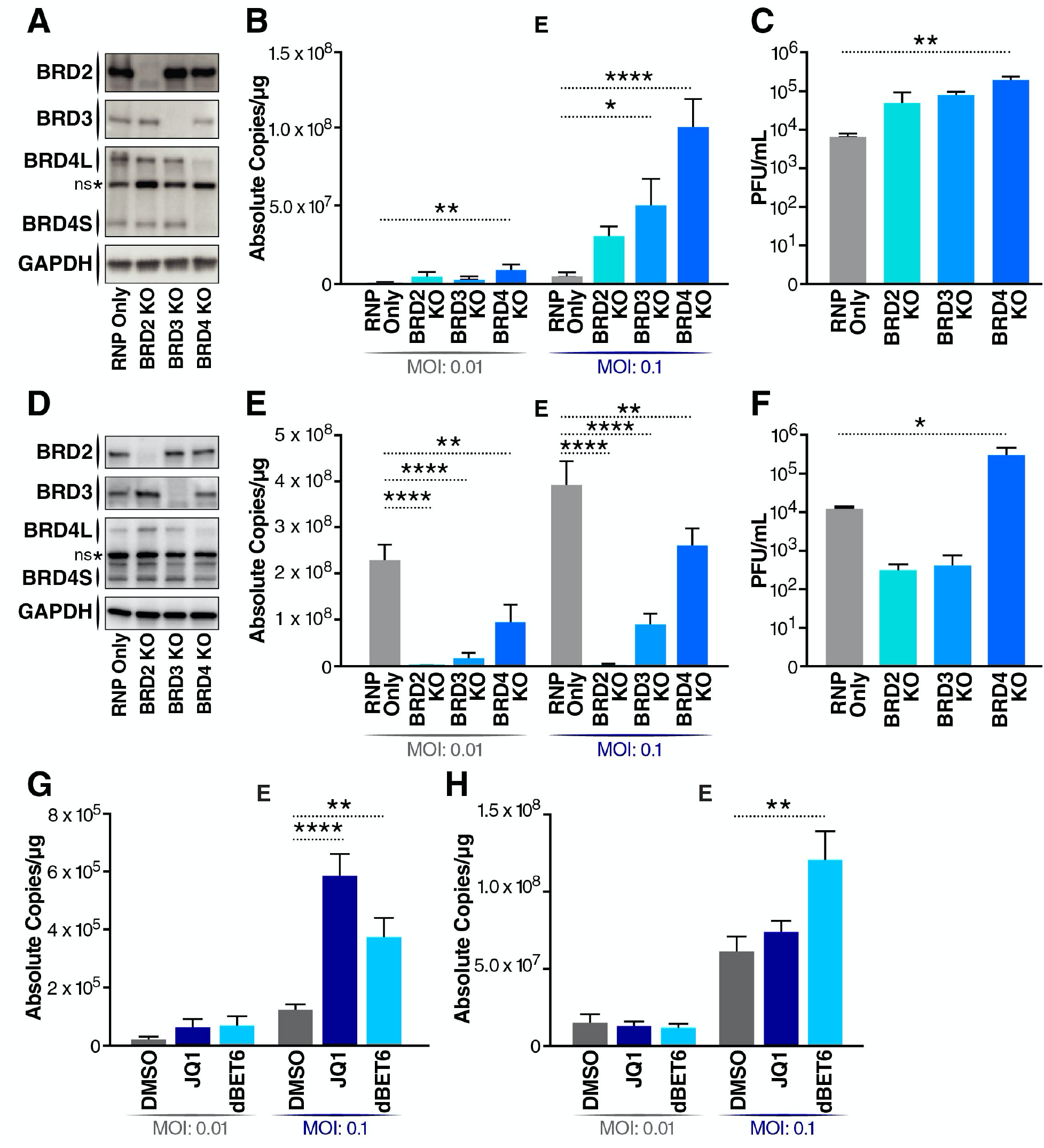
BRD3 and BRD4 proteins antagonize SARS-CoV-2 infection. A. Representative western blots from A549-ACE2 cells with indicated BET protein KOs. Lysates were probed for the epitope indicated beside each panel. *ns denotes a non-specific band. B. RT-qPCR of SARS-CoV-2 E RNA isolated 48 hours post infection (h.p.i) from A549-ACE2 cells with indicated BET protein KOs infected with SARS-CoV-2 (MOI of 0.01 or 0.1). Data are expressed in absolute copies/ug based on a standard curve of E gene with known copy number. Average of three independent experiments analyzed in triplicate ± SEM are shown and compared to RNP Only samples by ANOVA: ****p<0.0001, **p=0.0026, *p=0.0397. C. Plaque assay titers from supernatant of infected A549-ACE2 cells with indicated BET protein KOs at MOI of 0.1. Average of three independent experiments analyzed in duplicate ± SEM are shown and compared to RNP Only condition by ANOVA: **p=0.0049. D. Representative western blots from Calu3 cells with indicated BET protein KOs. Lysates were probed for the epitope indicated beside each panel. *ns denotes a non-specific band. E. RT-qPCR of SARS-CoV-2 E RNA isolated 48 h.p.i from Calu3 cells with indicated BET protein KOs infected with SARS-CoV-2 (MOI of 0.01 or 0.1). Data are expressed in absolute copies/ug based on a standard curve of E gene with known copy number. Average of three independent experiments analyzed in triplicate ± SEM are shown and compared to RNP Only samples by ANOVA: ****p<0.0001, **p<0.005. F. Plaque assay titers from supernatant of infected Calu3 cells with indicated BET protein KOs at MOI of 0.1. Average of three independent experiments analyzed in duplicate ± SEM are shown and compared to RNP Only condition by ANOVA: *p=0.0279. G. RT-qPCR of SARS-CoV-2 E RNA isolated 48 h.p.i. from A549-ACE2 cells infected with SARS-CoV-2 (MOI of 0.01 or 0.1) and concurrently treated with JQ1 (500nM) and dBET6 (500nM). Data are expressed in absolute copies/ug based on a standard curve of E gene with known copy number. Average of three independent experiments analyzed in triplicate ± SEM are shown and compared to DMSO condition by ANOVA: ****p<0.0001, **p<0.0069. H. RT-qPCR of SARS-CoV-2 E RNA isolated 48 h.p.i. from Calu3 cells infected with SARS-CoV-2 (MOI of 0.01 or 0.1) and treated with JQ1 (500nM) and dBET6 (500nM). Data are expressed in absolute copies/ug based on a standard curve of E gene with known copy number. Average of three independent experiments analyzed in triplicate ± SEM are shown and compared to DMSO condition by ANOVA: **p=0.0089.

In parallel, we generated BET protein KOs in Calu3 cells, airway epithelial cells with sufficient endogenous ACE2 expression to support SARS-CoV-2 infection. The knockouts were validated by western blotting for BET protein expression and infected with SARS-CoV-2 (Figure 1D). As expected, BRD2 KO decreased ACE2 transcript levels by ~80%, resulting in undetectable levels of viral RNA and >1 log decrease in infectious viral titers (Figure 1E, 1F, and S1A). BRD3 KO also reduced ACE2 expression by ~50% and viral replication about 3-fold (Figure 1E and S1A). Overall, BRD4 KO had the least effect on viral RNA expression and in fact increased infectious particle production despite decreasing ACE2 expression (Figure 1E, 1F, and S1A). These results show distinct effects of BET proteins on pre- and post-entry steps of SARS-CoV-2 infection and underscore BRD4’s role regulating the antiviral state after viral entry.

Next, we tested the pan-BET inhibitors, JQ1 and dBET6 that disrupt both BD1- and BD2-mediated interactions, in both A549-ACE2 and Calu3 cells. Viral RNA levels in A549-ACE2 cells significantly increased when JQ1 or dBET6 were added at the time of infection (Figure 1G). The same increase in viral RNA was observed with dBET6 treated Calu3 cells which express ACE2 endogenously (Figure 1H). This observation is in contrast to previous reports of prophylactic treatment of cell lines with the same compounds where ACE2 expression was reduced and viral entry was inhibited (Mills et al., 2021; Qiao et al., 2020; Tian et al., 2021). The use of the BET inhibitors at these effective concentrations was non-toxic in A549-ACE2 and Calu3 cells (Figure S1B and S1C). Collectively, these results establish that all BET proteins positively regulate ACE2 expression where BRD2 has the strongest effect, followed by BRD3 and BRD4. Based on the increases in viral RNA and infectious particle production we observed in BRD3 and BRD4 KO cells, we hypothesize these proteins are post-entry antagonists of SARS-CoV-2 viral replication; inactivating them, either genetically or chemically, exacerbates SARS-CoV-2 infection.

### Loss of BET Proteins Reduces Interferon and Proinflammatory Cytokine Expression

To test whether the loss of BET proteins reduces interferon production during SARS-CoV-2 infection, we infected Calu3 cells and then immediately treated them with either JQ1 or dBET6 for 48 hours, and analyzed mRNA expression of interferon-β (*IFNB1*), interferon stimulated gene 15 (*ISG15*) as well as of the proinflammatory cytokine interleukin 6 (*IL6*). We observed robust inductions of all three genes in control (DMSO-treated) cells that correlated with increasing MOI (Figure 2A). In contrast, upon JQ1 or dBET6 treatment, the expression of all three genes (*IFNB1*, *ISG15*, and *IL6*) was suppressed.

**Fig 2.**
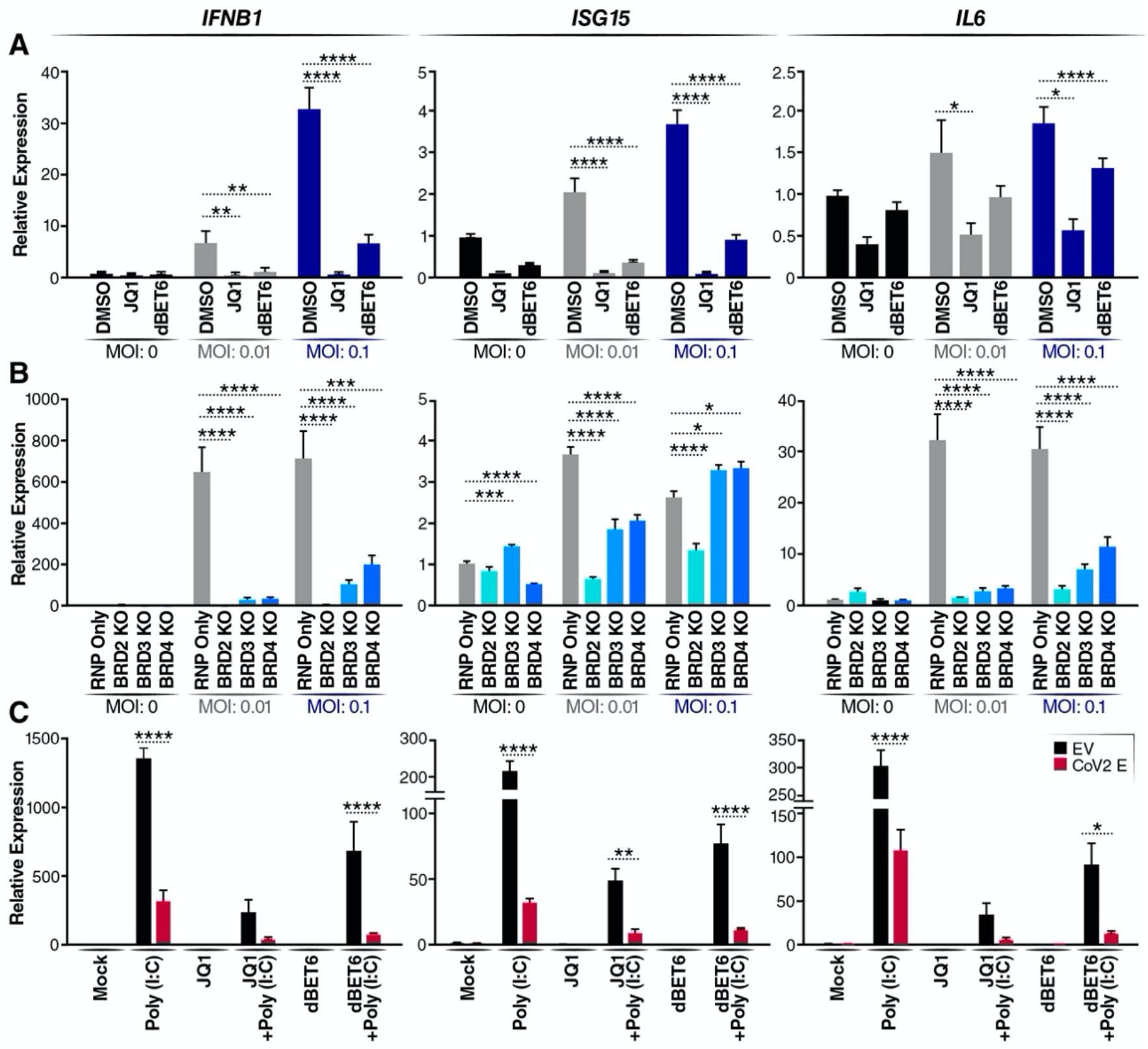
Loss of BET proteins reduces interferon and proinflammatory cytokine expression. A. RT-qPCR of RNA isolated 48 h.p.i. from Calu3 cells infected SARS-CoV-2 (MOI of 0.01 or 0.1) and concurrently treated with JQ1 (500nM) or dBET6 (500nM). Data are expressed relative to DMSO-treated cells for each respective MOI. Average of three independent experiments analyzed in triplicate ± SEM are shown and compared to DMSO condition by ANOVA for each MOI: ****p<0.0001, **p<0.005, *p<0.05. B. RT-qPCR of RNA isolated 48 h.p.i. from Calu3 cells with indicated BET protein KOs infected with SARS-CoV-2 (MOI of 0.01 or 0.1). Data are expressed relative to RNP Only cells for each respective MOI. Average of three independent experiments analyzed in triplicate ± SEM are shown and compared to RNP Only samples by ANOVA for each MOI: ****p<0.0001,***p<0.001 **p<0.005, *p<0.05. C. RT-qPCR of RNA isolated from A549 cells transfected with empty vector (EV) or SARS-CoV-2 E (CoV2 E) plasmid and treated with 10ng/ml Poly (I:C) for 24 hours with and without JQ1 (500nM) or dBET6 (500nM). Data are expressed relative to the mock treated cells. Average of three independent experiments analyzed in triplicate ± SEM are shown with ANOVA compared to mock: ****p<0.0001, **p=0.0097, *p=0.0139.

Next, we used the Calu3 BET KO cells to determine whether the loss of individual BET proteins was sufficient to suppress the induction of these genes. After infection with SARS-CoV-2, the control (RNP only) cells showed a robust induction of *IFNB1*, *ISG15*, and *IL6* at all MOIs (Figure 2B). In contrast, KO of all BET proteins significantly decreased expression of *IFNB1*, *ISG15*, and *IL6* at low MOI infection, phenocopying the effect of JQ1 and dBET6 on infected Calu3 cells (Figure 2B). Interestingly, at higher MOI, BET KO still suppressed *IFNB1* and *IL6* induction but *ISG15* expression was not significantly suppressed, implicating other ISGs as possible targets of BET action (Figure 2B). Notably, BRD2 KO cells showed the lowest interferon response reflecting their reduced infection due to low ACE2 levels (Figure S1A). These results underscore the role of different BET proteins in activating interferon and proinflammatory responses.

### SARS-CoV-2 E Protein Suppresses Host Antiviral Responses and Localizes to the Nucleus

The recent SARS-CoV-2 protein:protein interactome identified BRD2 and BRD4 as high confidence interactors with the viral E protein; BRD3 was also detected but just below the MiST scoring threshold (Figure 3A) (Gordon et al., 2020). We therefore tested whether E protein phenocopies the effect of BET inactivation. An E protein expression construct or the empty control vector (EV) was transfected into A549 cells and stimulated with poly (I:C) to induce interferon production (Figure 2C). Expression of E protein, similar to JQ1 and dBET6 treatment alone, dampened poly (I:C)-mediated induction of *IFNB1*, *ISG15*, and *IL6*, supporting the model that E protein evolved to suppress the antiviral response by inhibiting BET proteins.

**Fig 3.**
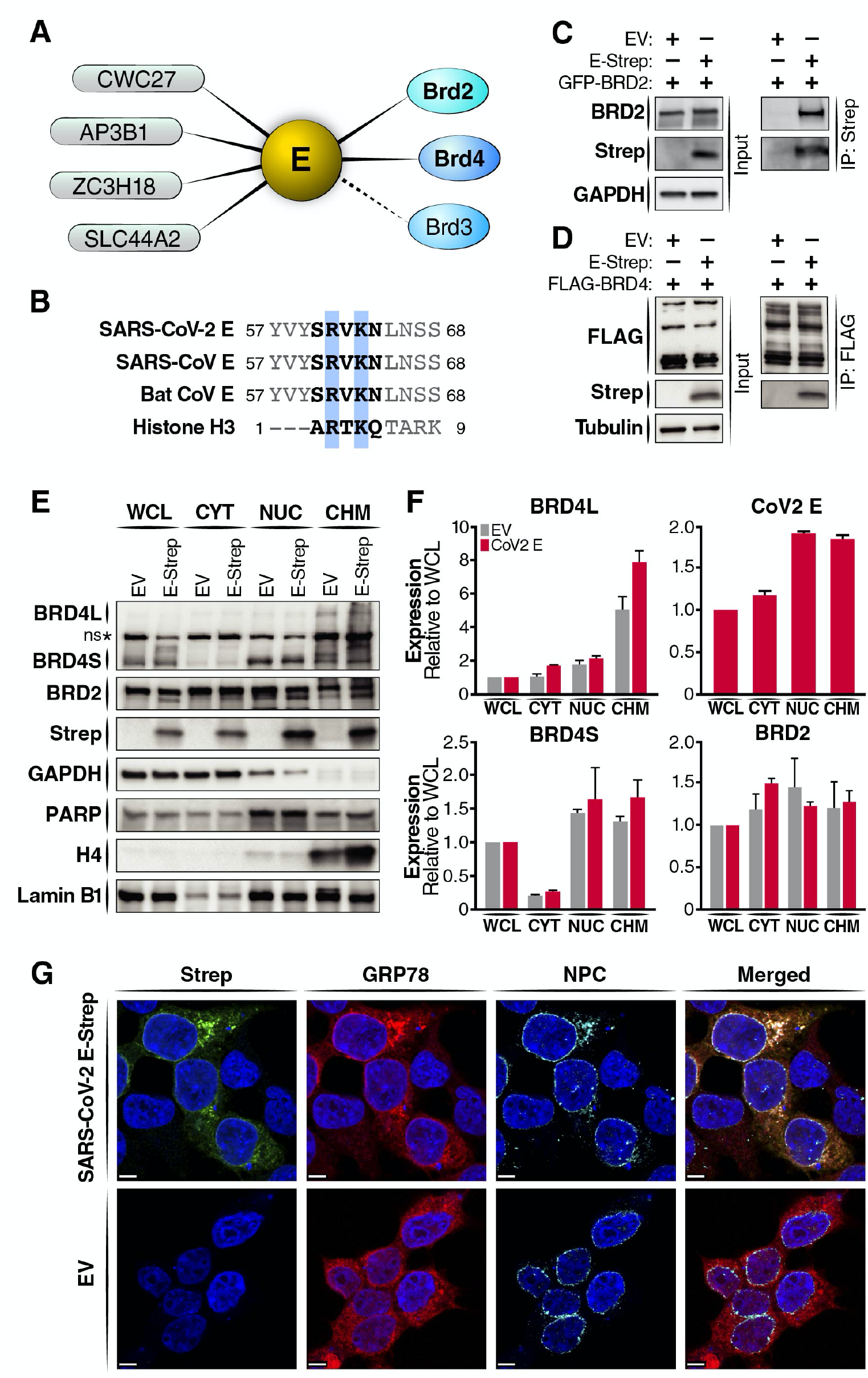
SARS-CoV-2 E protein interacts with BET proteins at the nuclear periphery. A. Host protein interactors, including members of the BET protein family (BRD2, BRD3, BRD4), of the SARS-CoV-2 E protein (Gordon et al. 2020.). High confidence interactors above the MiST threshold are shown in solid lines and interactors below the threshold are shown with dashed lines. B. The E protein sequences from human and bat coronaviruses share a histone H3-like motif. SARS-CoV (YP_009724392.1), SARS-CoV-2 (NP_828854.1), Bat CoV (AGZ48809.1), and histone H3 (P68431). C. Immunoprecipitation of overexpressed Strep-tagged SARS-CoV-2 E (E-Strep) protein from transfected HEK293T whole-cell lysates, followed by western blotting using Strep, BRD2, and GAPDH antibodies. EV is empty vector control. D. Immunoprecipitation of overexpressed FLAG-tagged BRD4 from transfected HEK293T whole-cell lysates, followed by western blotting using FLAG, Strep, and tubulin antibodies. EV is empty vector control. E. Western blotting of whole-cell lysate (WCL), cytoplasmic (CYT), nuclear (NUC), and chromatin (CHM) fractions from HEK293T cells transfected with EV (empty vector) or Strep-tagged SARS-CoV-2 E (Strep-E) with indicated antibodies. *ns denotes a non-specific band. F. Quantification of BRD2, BRD4L, BRD4S, and SARS-CoV-2 E (Strep-E) band intensities in the CYT, NUC, and CHM fractions expressed relative to WCL. Average of two biological replicates is shown. G. Representative confocal microscopy images of HEK293T cells transduced with SARS-CoV-2 E or control (empty vector). Cells were processed for immunostaining with Strep (SARS-CoV-2 E, green), GRP78 (ER, red), NPC (nuclear pore complexes, turquoise), and Hoechst (nuclei, blue). Scale bars, 5um.

We previously proposed that a short motif in E protein’s C-terminus mimics the N-terminus of histone H3 (Figure 3B). This motif was conserved in the E proteins of other coronaviruses, supporting a broader model where coronavirus E proteins may neutralize the antiviral response by antagonizing BET proteins. Mechanistically, this may work by inhibiting the interaction of BET proteins with chromatin. To test this model, we performed co-immunoprecipitation studies in HEK293T cells overexpressing tagged forms of E and BET proteins. We found that SARS-CoV-2 E protein (Streptavidin (Strep)-tagged) interacted with BRD2 (green fluorescent protein (GFP)-tagged) and BRD4 (FLAG-tagged) (Figure 3C and 3D). Notably, E protein interacted with both isoforms of BRD4, pointing to the N-terminus with conserved BDs and ET domain as the interaction site (Figure S2A and S2B). Next, we performed cellular fractionation studies in HEK293T cells overexpressing Strep-tagged E protein. Half of total E was enriched in the nuclear fraction, and the majority of this fraction co-fractionated with chromatin (Figure 3E and 3F). These fractions also contained the majority of BET proteins. We did not observe a difference in BET protein fractionation between cells expressing E protein or the empty vector, excluding the possibility that E protein grossly mislocalizes BET proteins. This was further confirmed by confocal microscopy, where E protein, a membrane-bound protein, showed a ring-like perinuclear localization that overlapped with staining of the cellular nuclear pore complex (NPC) (Figure 3F) (Schoeman and Fielding, 2019). Of note, previous reports of E proteins from SARS-CoV (SARS) and MERS-CoV (MERS) imaged in infected cells also showed a perinuclear localization (Nal et al., 2005; Nieto-Torres et al., 2011). However, reliable antibodies to detect untagged SARS-CoV-2 E protein during infection are still missing, making staining of endogenous E during infection technically impossible. Overall, our data support a model where the E:BET protein interaction occurs on chromatin in the nuclear periphery.

### Acetylated SARS-CoV-2 E Protein Binds the Second Bromodomain of BRD4

Next, we tested whether E protein is acetylated in cells in order to act as a histone mimetic and interact with BDs. For this, we immunoprecipitated FLAG-tagged E protein from transfected HEK293T cells and probed for acetylation with pan-acetyl-lysine antibodies. We found that E protein is acetylated in cells (Figure 4A). The acetyl-lysine signal was further enhanced when cells were treated with a pan-inhibitory cocktail of cellular histone deacetylase (HDAC) inhibitors, showing that E protein undergoes reversible acetylation in host cells (Figure 4A).

**Fig 4.**
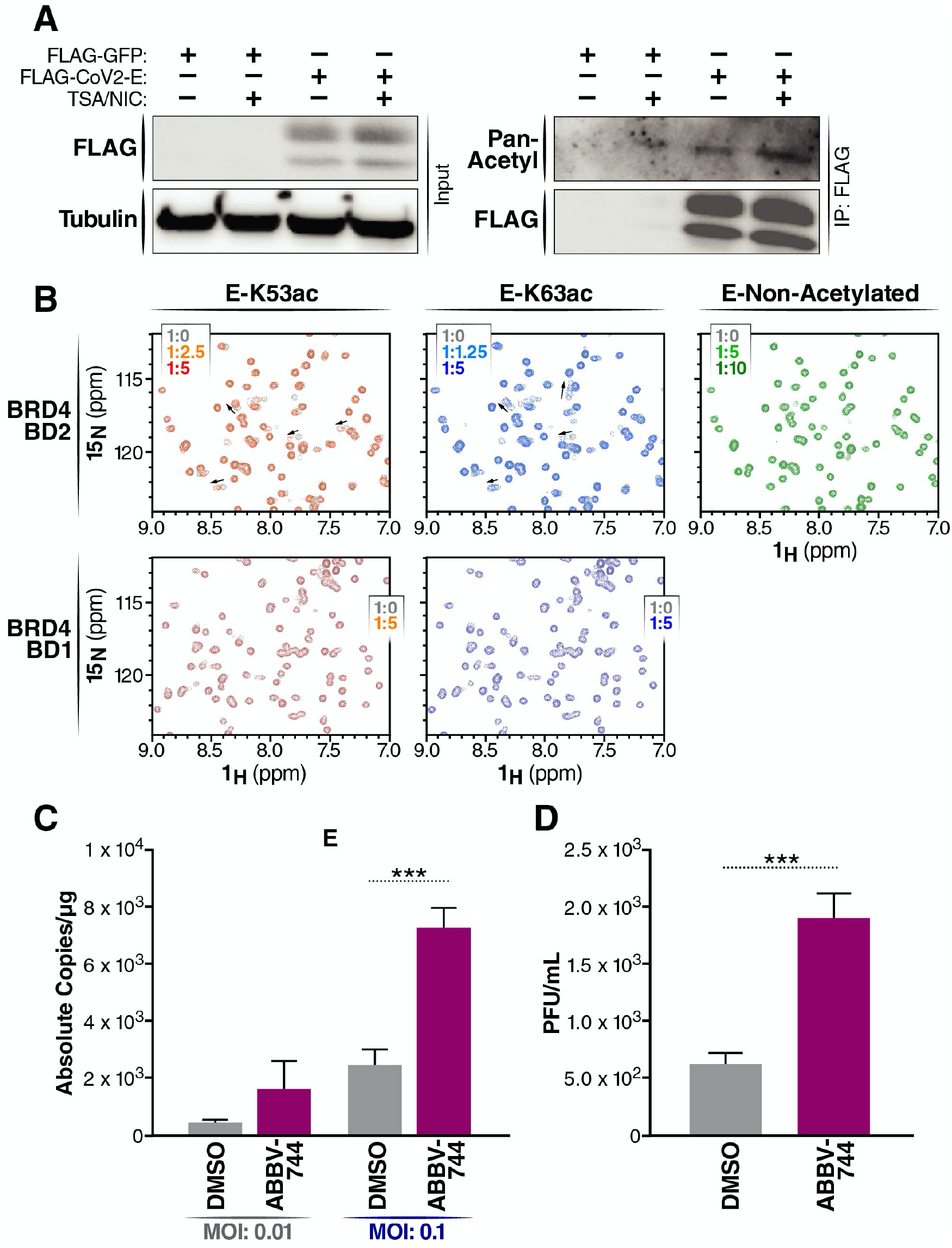
Acetylated SARS-CoV-2 E protein binds the second bromodomain of BRD4. A. Immunoprecipitation of overexpressed FLAG-tagged SARS-CoV-2 E protein (FLAG-E) from transduced HEK293T whole-cell lysates with and without TSA (1uM) and NIC (1mM), followed by western blotting using FLAG, pan acetyl lysine, and tubulin antibodies. FLAG-GFP is vector control. B. 2D ^1^H ^15^N HSQC spectra measured after addition of either E - K53ac (residues 48-58, acetylated), E – K63ac (residues 58-68, acetylated) or Non-Acetylated (residues 58-70) peptides to ^15^N labelled BD1 or BD2. Arrows indicate chemical shift perturbations of peaks. Protein-to-ligand ratio is indicated. C. RT-qPCR of SARS-CoV-2 E RNA isolated 48 h.p.i. from A549-ACE2 cells infected with SARS-CoV-2 (MOI of 0.01 or 0.1) and concurrently treated with BD2-selective inhibitor, ABBV-744 (500nM). Data are expressed in absolute copies/ug based on a standard curve of E gene with known copy number. Average of three independent experiments analyzed in triplicate ± SEM are shown and compared to DMSO by Students t-test: ***p=0.0004. D. Plaque assay titers from supernatant of infected A549-ACE2 cells at MOI of 0.1 treated with ABBV-744 (500nM). Average of three independent experiments analyzed in duplicate ± SEM are shown and compared to DMSO by Student’s t-test: ***p=0.0001.

We used nuclear magnetic resonance (NMR) titration experiments to assess binding of E peptides spanning the histone-like motif to recombinant BD1 and BD2 portions of BRD4. The histone motif contains two lysines, K53 and K63, and we used peptides that were acetylated at either position or were non-acetylated. In two-dimensional ^1^H-^15^N heteronuclear single quantum coherence (HSQC) spectra of BD2, we saw significant chemical shift perturbations of peaks upon the addition of either E-K53ac or E-K63ac peptides, indicating BD2 is able to bind either acetyl-lysine position (Figure 4B, upper panels). BD2 was found to have a higher affinity for the K63ac peptide (*K*_d_ value of 173 ± 57 μM) than for the E K53ac peptide (*K*_d_ value of 610 ± 162 μM) (Figure S3A). The interaction was acetylation-dependent as the addition of non-acetylated E peptide did not elicit changes in chemical shifts in BD2 as observed with the acetylated peptides. In order to determine the peptide binding site, the backbone amides most perturbed upon peptide binding were mapped onto the surface of BD2. Both peptides were found to occupy a similar binding site to where acetylated lysine peptides have been shown to bind (Figure S3B and S3C). Unlike with BD2, little to no chemical shift perturbations in HSQC spectra were observed upon addition of the acetylated E peptides to BD1 indicating very weak to no interaction of acetylated E protein with BD1 (Figure 4B, lower panels).

Overall, these data show that the E protein is acetylated and that the interaction of BRD4 with acetylated E protein involves the second bromodomain of BRD4. The relevance of the second bromodomain to the antiviral function of BRD4 was further explored in studies with ABBV-744, a BET inhibitor that specifically targets BD2-mediated interactions (Faivre et al., 2020; Sheppard et al., 2020). In A549-ACE2 cells treated with ABBV-744 at the time of infection, SARS-CoV-2 viral RNA and infectious particle production were significantly increased (3-fold for both) when compared to cells treated with the vehicle alone, demonstrating that selective inhibition of BD2 alone is sufficient to suppress the antiviral function of BET proteins (Figure 4C and 4D).

### BET Inhibitors Enhance SARS-CoV-2 Infection in K18-hACE2 mice

To test the post-entry effects of BET inhibitors, JQ1 and ABBV-744, *in vivo*, we used K18-hACE2 transgenic mice constitutively expressing the human ACE2 receptor in epithelial tissues from the cytokeratin 18 (K18) promoter (Jia et al., 2005). Mice were infected intranasally, and BET inhibitors were administered daily intraperitoneally or orally starting on the day of infection (Figure 5A). DMSO at the vehicle concentration was used as a control.

**Figure 5.**
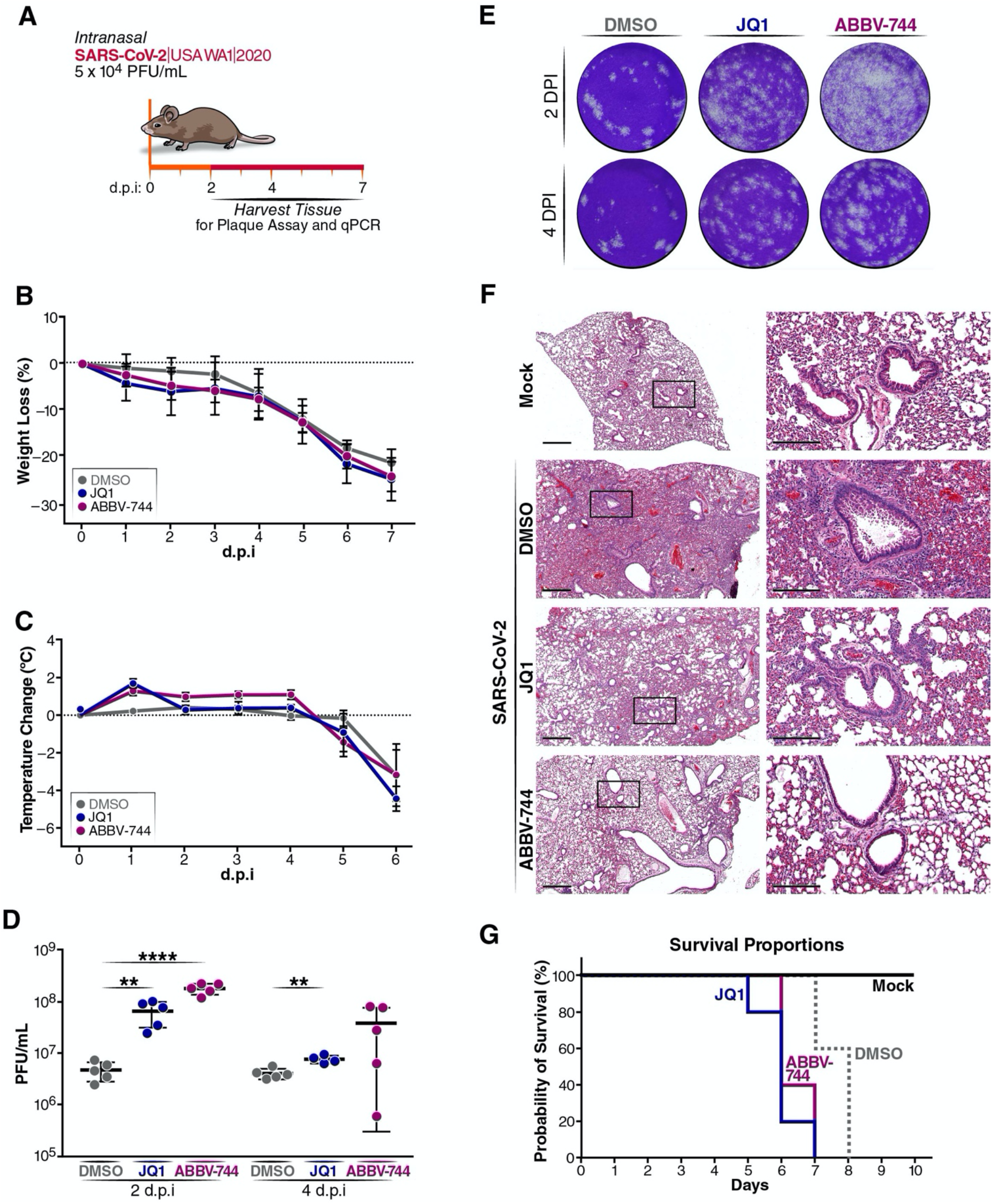
BET inhibitors enhance SARS-CoV-2 infection in K18-hACE2 mice. A. Schematic of the experiment. B. Severe weight loss of SARS-CoV-2 infected mice due to severe lung infection that required euthanasia. n=5 mice per group. C. Changes in body temperature of SARS-CoV-2 infected mice. n=15 mice per group. D. Plaque assay titers from the lungs of infected mice. Average of 5 mice per group were analyzed, average ± SD are shown and compared to DMSO by Student’s t-test. 2 dpi: **p=0.0041, ****p<0.0001. 4dpi: **p=0.0021. E. Representative images of plaque assays at the same dilution from infected mice at 2 and 4 days post infection. F. Representative images of &E staining of lung tissue at 7 days post infection. Left panels, scale bars, 600um. Right panels enlarged images of left panel, scale bars, 200um. G. Survival curve of mock (not infected) and infected DMSO-(n=15), JQ1-(n=15), and ABBV-744-(n=15) treated mice.

As expected, all infected mice exhibited weight loss and change in body temperatures as a sign of infection (Figure 5B and 5C). Severe disease was observed in the BET inhibitor treated mice, where all JQ1- and ABBV-744-treated mice exhibited severe weight loss (>20%) and were hunched and lethargic on day 5 and 6 post infection, while the DMSO-treated group reached to same phenotype at day 6-8 post infection. Viral replication in all treatment groups was analyzed at day 2 and 4 post infection. During these time points, we observed increased viral titer in the lungs of JQ1-and ABBV-744-treated mice (Figure 5D and 5E). Specifically, there was an 18-fold increase in infectious particle production in JQ1-treated mice and a 37-fold increase in ABBV-744-treated mice compared to DMSO at day 2 post infection. Similar to the *in vitro* treatment, the JQ1- and ABBV-744-treated mice experienced significant decreases in the expression of interferons, cytokines, and chemokines, including *ISG15*, *IL6*, *RIG-I*, *IL1a*, *CCL2*, *CXCL9*, and *CXCL10* (Figure S4A). The decrease in these host defense gene products paired with the subsequent increase in infectious particle production was accompanied by a decrease in immune infiltration in the lung at 7 days post infection in the JQ1- and ABBV-744-treated mice compared to DMSO (Figure 5F). The DMSO treatment showed widespread large inflammatory foci consistent with interstitial pneumonia with perivascular inflammation comprised of lymphocytes, macrophages and neutrophils.

Alongside disease severity in BET inhibitor-treated mice was characterized by extreme hyperthermic and hypothermic conditions compared to DMSO treatment groups (Figure 5C). Furthermore, during dissection of the mice, we observed increased gut inflammation in JQ1 (40%) and ABBV-744 (60%) compared to DMSO (20%) at day 4 post infection (Figure S4B). All animals in the JQ1 and ABBV-744 treatment groups showed the hunched-back phenotype at day 5 and 6 post infection which is earlier compared to DMSO treatment (Figure S4C). 60-80% animals with BET inhibitor treatment reached humane endpoint at day 6 post infection whereas 100% of the animals in the DMSO-treated group survived at this time (Figure 5G). These data underscore the finding that BET proteins regulate an antiviral state during viral infection that prevent exacerbation of virally induced disease and document the clinical relevance of the second bromodomain of BET proteins, also targeted by the viral E protein.

## DISCUSSION

We report that BET proteins (BRD4 > BRD3 > BRD2) block SARS-CoV-2 infection at the post-entry level as each of them is necessary for full induction of the type I interferon response and the IL-6 proinflammatory cytokine. Viral replication is exacerbated after chemical or genetic inactivation of BET proteins, underscoring the critical role of this antiviral step in controlling SARS-CoV-2 infection. We further report a new function of the SARS-CoV-2 E protein in antagonizing interferon and ISG expression. E protein is acetylated in cells and in an acetylated form binds to the second bromodomain of BRD4. Lastly, BET inhibitor treatment increases viral titers in the lungs of SARS-CoV-2 infected mice and results in higher morality. As the SARS-CoV-2 E protein is functionally similar to BET antagonists in suppressing the interferon response and directly binds one of the two bromodomains, we propose a model where E protein evolved to neutralize the antiviral gene activation mediated by BET proteins to promote efficient SARS-CoV-2 replication, a process further enhanced with BET inhibitor treatment.

Our data highlight the unique and overlapping roles of BET proteins during SARS-CoV-2 infection where individual BET proteins may serve proviral and antiviral roles. We show that BET proteins can promote or block SARS-CoV-2 infection, and this is likely dependent on the specific gene targets with which individual BET proteins interact.

Among the BET proteins, BRD2 is the most proviral because it positively regulates the expression of the SARS-CoV-2 entry receptor ACE2. In contrast, BRD4 has the strongest antiviral function due to its co-activator role in the induction of interferon genes. Differences between BRD2 and BRD4 lie in their domain structure; while sharing roughly 70% sequence similarity in their N-terminus including the tandem BDs, the C-terminus of BRD4L contains the PID which enhances the recruitment of P-TEFb to BET target genes (Bisgrove et al., 2007; Sheppard et al., 2020). Small molecule BET inhibitors reduce the recruitment of BET proteins, along with P-TEFb, to interferon target genes upon interferon-β or Toll-like receptor ligand stimulation (Gilan et al., 2020; Malik et al., 2015; Patel et al., 2013). Furthermore, BRD4 plays a critical role in coordinating both positive and negative regulation of paused RNA polymerase II at the transcription start sites of ISGs by recruiting NELF/DSIF (negative elongation factor/DRB sensitivity-inducing factor) to fine-tune ISG expression (Patel et al., 2013). On the other hand, BRD2 functions as a chromatin organizer that assembles enhancer elements (Cheung et al., 2017). The depletion of BRD2 versus BRD4 results in distinct transcriptional changes, suggesting non-overlapping roles in function (Andrieu and Denis, 2018; Deeney et al., 2016). In contrast, the identification of a conserved motif B in BET proteins that facilitates heterodimerization of proteins within this family may explain partial functional redundancy and shared chromatin occupancy of BRD2 and BRD4 (Garcia-Gutierrez et al., 2012). The fact that BRD2 and BRD4 play opposing roles in the regulation of SARS-CoV-2 highlights the complexity of genes regulated by BET proteins and future studies aimed at disentangling these regulatory networks will be of value to the field.

We also show that SARS-CoV-2 E protein targets the second bromodomain of BRD4. Both BD1 and BD2 share the characteristic four α-helix bundle structure of bromodomains (Zeng and Zhou, 2002). However, BD1 has a partly basic surface and BD2 a region of acidic residues proximal to the binding pockets of these two bromodomains for acetylated lysines; this difference may explain their recognition of acetylated lysine motifs in different sequence contexts (Vollmuth et al., 2009). Research has classically focused on histones as BD targets, but BD2 of BRD4 also binds the acetylated cyclin T1 subunit of P-TEFb (Schroder et al., 2012). Recent studies defining the unique roles of BD1 and BD2 in disease models indicated that inhibiting BD2 preferentially blocks stimulus-mediated induction of gene expression programs without affecting pre-existing transcriptional programs (Gilan et al., 2020). BD1-specific inhibitors are effective in inhibiting cell proliferation in cancer models, whereas BD2-specific inhibitors had the greatest effect in ameliorating inflammatory and autoimmune diseases (Gilan et al., 2020). Our finding that SARS-CoV-2 E protein binds BD2 of BRD4 supports the notion that BD2 is important for rapid gene expression, especially in inflammatory disease settings such as COVID-19.

SARS-CoV-2 E protein joins a growing number of viral proteins containing histone-like motifs in their sequences. Histone tails are often mimicked by viruses since many host chromatin factors interact with histones to modulate gene expression (Schaefer et al., 2013). A prominent example of a viral histone mimetic is NSP1 of influenza A H3N2 subtype, which contains a short histone H3-like motif capable of sequestering the transcription elongation factor, PAF1, to prevent the expression of antiviral genes (Qin et al., 2014). Similarly, Human Immunodeficiency Virus (HIV-1) factor Tat is reversibly acetylated at multiple lysines in its basic domain to recruit key host transcriptional regulators in a coordinated manner, enhancing viral transcription (Dorr et al., 2002; Huo et al., 2011; Kiernan et al., 1999; Ott et al., 1999; Pagans et al., 2005). A recent study identified the capsid protein of the Yellow Fever Virus as a histone H4 mimetic with acetylated lysine residues that bind BRD4 to interfere with gene regulation (Mourao et al., 2020). Overall, BRD4 appears to be a hotspot of viral antagonism due to its critical role in positively regulating the antiviral response.

The E proteins of coronaviruses are similar in domain structure: a short, hydrophilic N-terminus, followed by a hydrophobic transmembrane domain, and a subsequent C-terminus that comprises the majority of the protein and contains a PDZ-binding motif (PBM). Among the E proteins of human pathogenetic coronaviruses, those from SARS and SARS-CoV-2 share ~95% sequence similarity along with the histone H3-like motif whereas the MERS E protein shares only ~35% similarity to SARS and SARS-CoV-2 E proteins and does not have the histone-like motif (Schoeman and Fielding, 2020). The histone-like motif of interest is also present in common coronavirus NL63 and bat coronavirus SHC014, suggesting that the function of E:BET proteins could also be conserved in these viruses. Future work should address if E proteins from these coronaviruses are similarly capable of derailing antiviral responses through BRD4 antagonism.

SARS-CoV-2 E protein is only one of several factors encoded by this virus capable of suppressing host IFN-I responses. Other anti-interferon factors are: NSP6 and NSP13 which inhibit TANK binding kinase 1 (TBK1) and interferon-regulatory factor 3 (IRF3) activation; ORF6 which prevents IRF3 nuclear translocation and suppresses ISG promoter activation; ORF9b which suppresses retinoic acid-inducible gene I-mitochondrial antiviral-signaling protein (RIG-I-MAVS) signaling and NSP1, NSP6, NSP13, ORF3a, ORF7a/b, and membrane (M) protein which block signal transducer and activator of transcription 1/2 (STAT1/2) phosphorylation and STAT1 nuclear translocation (Lei et al., 2020; Wu et al., 2021; Xia et al., 2020). The fact that SARS-CoV-2 targets the interferon response at these different levels emphasizes the importance of overcoming this innate immune pathway to successfully establish and maintain viral infection.

Since the beginning of the pandemic, there has been growing interest in the prophylactic application of BET inhibitors to prevent SARS-CoV-2 infection as they reduce ACE2 expression. All previous studies relied on multi-day pretreatment of cell lines, organoids, and primary cells with BET inhibitors to inhibit infection (Gilham et al., 2021; Mills et al., 2021; Qiao et al., 2020; Tian et al., 2021). In contrast, our study focuses on the concurrent administration of BET inhibitors, JQ1 and ABBV-744, at the time of infection. Similar to our observations, Mills et al. found that therapeutic application of K18-hACE2 mice with BET inhibitor, INCB054329, resulted in severe lung pathology and significant viral RNA in the lung. We demonstrate the therapeutic application of the BET inhibitors worsens viral pathogenesis as evidenced by hypothermic and hyperthermic conditions, severe weight loss, and inflammation of gut, underscoring the clinical significance of post entry restriction of SARS-CoV-2 by BET proteins.

Our study sheds new light on the antiviral function of BET proteins during SARS-CoV-2 infection, highlighting their contrasting functions at different stages of the viral life cycle. Whether SARS-CoV-2 evolved the anti-BET protein function of the E protein because of the unique influence of BRD2 on ACE2 receptor levels or the necessity for a more surgical inactivation of BET proteins as transcriptional coactivators necessary for antagonizing the virus remains to be determined. Because most BET inhibitors do not distinguish between the different BET proteins and inhibit all of them indiscriminately, our study urges caution with the clinical use, prophylactic or therapeutic, of pan-BET inhibitors in people at risk or afflicted with COVID-19.

## RESOURCE AVAILABILITY

### Lead contact

Further information and requests for resources and reagents should be directed to and will be fulfilled by the Lead Contact, Melanie Ott.

### Materials availability

This study did not generate new unique reagents.

### Data and code availability

This study did not generate/analyze datasets or code.

## EXPERIMENTAL MODEL AND SUBJECT DETAILS

### Mammalian cell lines and culture conditions

HEK293T and Vero-E6 were obtained from ATCC were cultured in DMEM (Corning) supplemented with 10% fetal bovine serum (FBS) (GeminiBio), 1% glutamine (Corning), and 1% penicillin-streptomycin (Corning) at 37°C, 5% CO_2_.

Calu3 cells were obtained from ATCC and cultured in AdvancedMEM (Gibco) supplemented with 2.5% FBS, 1% GlutaMax, and 1% penicillin-streptomycin at 37°C, 5% CO_2_.

A549 cells stably expressing ACE2 (A549-ACE2) were a gift from O. Schwartz. A549-ACE2 cells were cultured in DMEM supplemented with 10% FBS, blasticidin (20 μg/ml) (Sigma) and maintained at 37°C with 5% CO_2_. Short Terminal Repeat (STR) analysis by the Berkeley Cell Culture Facility on 17 July 2020 authenticates these as A549 cells with 100% probability.

### Generation of CRISPR A549-ACE2 and Calu3 KO cell lines

sgRNAs were designed according to Synthego’s multi-guide gene knockout. Briefly, three sgRNAs are bioinformatically designed to work in a cooperative manner to generate small fragment deletions in early exons causing knock out. These fragment deletions are larger than standard indels generated from single guides. The genomic repair patterns from a multi-guide approach are highly predictable based on the spacing of and on design constraints that limit off-targets, resulting in a higher probability of protein knockout phenotype.

To induce gene knockout, 10 pmol Streptococcus Pyogenes NLS-Sp.Cas9-NLS (SpCas9) nuclease (Aldevron, 9212) was combined with 30 pmol total synthetic sgRNA (10 pmol each sgRNA) (Synthego) to form ribonucleoproteins (RNPs) in 20uL total volume with SE Buffer for Calu-3 and A549-ACE2 cells. The RNP assembly reaction was mixed by pipetting up and down and incubated at room temperature for 10 minutes. Cells were resuspended in transfection buffer according to cell type and added to the preformed RNP solution and gently mixed. Nucleofections were performed on a Lonza nucleofector system (Lonza, AAU-1001) using program CM-130 and CM-120 for Calu-3 and A549-ACE2 cells, respectively. Two days post-nucleofection, DNA was extracted with DNA QuickExtract (Lucigen, QE09050). Amplicons for indel analysis were generated by PCR using AmpliTaq Gold 360 polymerase (Thermo Fisher Scientific, 4398813). PCR products were cleaned-up and analyzed by Sanger sequencing. Sanger data files and sgRNA target sequences were input into Inference of CRISPR Edits (ICE) analysis (https://ice.synthego.com) to determine editing efficiency and to quantify generated indels (Hsiau T., 2019). A list of sgRNA sequences and genotyping primers can be found in Table S1.

### SARS-CoV-2 virus culture

SARS-CoV-2 Isolate USA-WA1/2020 (BEI NR-52281) was used for all infection studies. All live virus experiments were performed in a Biosafety Level 3 laboratory. SARS-CoV-2 stocks were propagated in Vero-E6 cells and their sequence verified by next-generation sequencing. Viral stock titer was calculated using plaque forming assays.

### Plasmids

The plasmids expressing the envelope protein of SARS-CoV-2 were generous gifts from Dr. Nevan Krogan (UCSF, The Gladstone Institutes). The majority of BRD4 plasmids were generous gifts from Dr. Eric Verdin (The Buck Institute for Research on Aging, Novato, CA). All plasmids and corresponding sequence information are available upon request.

### Cell Fractionation

Cell fractionation was performed with the NE-PER Nuclear and Cytoplasmic Extraction Kit (Thermo Fisher Scientific, 78835) with additional modifications to extract the chromatin fraction. Following the extraction of the nuclear fraction per manufacturer’s instructions, the resulting pellet was resuspended in NER with MNase (NEB, M0247S), Halt protease inhibitor cocktail (Thermo Fisher Scientific, 1861282), and CaCl_2_ (Sigma, C4901). Samples were vortexed on the highest setting and sheared on the Bioruptor (Diagenode) with 10 cycles of 10 seconds on and 10 seconds off on medium setting. The samples were pelleted by centrifugation and the resulting supernatant was the chromatin fraction.

### Western Blot Analysis

Cells were lysed in RIPA lysis buffer (50 mM Tris-HCl [pH 8], 150 mM NaCl, 1% NP-40, 0.5% sodium deoxycholate, 0.1% SDS, supplemented with Halt protease inhibitor cocktail) to obtain whole-cell lysates or lysed using the NE-PER nuclear and cytoplasmic extraction kit to obtain cytoplasmic, nuclear, and chromatin fractions. Protein concentrations of cell fractions were determined using a BCA Assay (Thermo Fisher Scientific, 23225), and normalized among samples per experiment before analysis via western blotting using standard techniques. Proteins were visualized by chemiluminescent detection with ECL on ChemiDoc MP (Bio-Rad). Antibodies: BRD2 (Abcam, ab139690), BRD3 (Abcam, ab50818), BRD4 (Abcam, ab128874), LaminB1 (Abcam, ab16048), PARP (Cell Signaling, 9532S), GAPDH (Cell Signaling, 5174S), Tubulin (Abcam, ab7291), Strep (Qiagen, 1023944), FLAG (Sigma, F3165), Histone H4 (Cell Signaling, 13919S), pan acetyl lysine (Cell Signaling 9441S; Abcam, ab80178), Rabbit IgG-HRP (Bethyl, A120-201P), and Mouse IgG-HRP (Bethyl, A90-516P).

### Quantitative Polymerase Chain Reaction

RNA was extracted from cells or supernatants using RNA STAT-60 (AMSBIO, CS-110) and the Direct-Zol RNA Miniprep Kit (Zymo Research, R2052). RNA from cells or supernatant was reverse-transcribed to cDNA with iScript cDNA Synthesis Kit (Bio-Rad, 1708890). qPCR reaction was performed with cDNA and SYBR Green Master Mix (Thermo Scientific) using the CFX384 Touch Real-Time PCR Detection System (Bio-Rad). See Table S1 for primers sequences. E gene standards were used to generate a standard curve for copy number quantification. E gene standard was generated by PCR using extracted genomic SARS-CoV-2 RNA as template. A single product was confirmed by gel electrophoresis and DNA was quantified by Nanodrop.

### Immunoprecipitation

Transfected HEK293T cells were lysed in IP buffer (50mM Tris-HCl, 150mM NaCl, 1mM EDTA, 1% Triton-X, supplemented with Halt protease inhibitor cocktail) and 1mg of lysate was incubated with FLAG-M2 magnetic beads (Sigma, M8823) or Strep-Tactin Sepharose resin (iba Life Science, 2-1201-002) overnight rotating at 4°C. Bound protein was washed five times with IP buffer and eluted by peptide competition (Sigma, F4799) or Strep-Tactin elution buffer (iba Life Sciences, 2-1000-025). Eluted samples were analyzed by western blot.

### Immunofluorescence Microscopy

Transfected HEK293T cells were plated onto rat tail collagen (EMS, 72295) coated, 22- by 22-mm no. 1.5 coverslips. Cells were fixed in 4% paraformaldehyde, permeabilized with methanol on ice for 10 minutes, and blocked in 3% bovine serum albumin. Cells were then immunostained with the indicated antibodies: STREP (Qiagen, 1023944), FLAG (Sigma, F3165), LaminB1-AlexaFluor647 (Abcam, ab16048), Hoescht 33342 (Invitrogen, H3570), Mouse IgG-AlexaFluor549 (Invitrogen, A11005), and Rabbit IgG-AlexaFluor488 (Invitrogen, A11008). Coverslips were mounted onto glass slides using ProLong Gold Antifade Mountant (Invitrogen, P36934) and analyzed by confocal microscopy (Olympus FV3000RS) using a Olympus UPLAN S-APO 60X OIL OBJ,NA1.35,WD0.15MM objective. The resulting Z-stack was reconstructed and rendered in 3D using Imaris software (Oxford Instruments).

### Expression and purification of Brd4 bromodomains

His-SUMO-BD1 (42-168) and BD2 (349-460) constructs were expressed in LOBSTR E. coli cells (Kerafast, EC1002). Expression and purification of both constructs followed the same protocol. For NMR studies the His-SUMO-BD constructs were expressed in M9 minimal media containing ^15^N ammonium chloride. Cells were induced with 0.1 mM IPTG and grown at 16 °C overnight before the pellet was collected. The cells were resuspended in lysis buffer (50 mM Tris, 200 mM NaCl, pH 7.5), lysed by sonication and centrifuged. The supernatant was purified using Ni resin equilibrated in binding buffer (50 mM Tris, 200 mM NaCl, 20 mM imidazole, pH 7.5), washed first with lysis buffer and 30 mM imidazole and then a final wash with lysis buffer and 50 mM imidazole. The protein was eluted from the Ni resin after incubation with SUMO protease Ulp1 in order to remove the His-SUMO tag. The cleaved proteins were then further purified by size-exclusion chromatography using a Hiload 26/60 Superdex 75 gel filtration column (GE, GE28-9893-34) in a buffer of 50 mM sodium phosphate, 100 mM NaCl pH 7.4 buffer before being concentrated and flash–frozen.

### NMR binding experiments

For 2D ^1^H, ^15^N HSQC peptide titration experiments, 50 μM ^15^N-labeled BD1 or BD2 in 50 mM sodium phosphate, 100 mM NaCl, 5 % D_2_O, pH 7.4 buffer was used, and spectra were measured after each addition of E protein peptide. The 2D ^15^N-FHSQC spectra for the E protein peptide titration series were recorded on a Bruker 500 MHz AVANCE DRX spectrometer equipped with an actively shielded Z-gradient QCI cryoprobe (^15^N/^13^C/^31^P, ^1^H) using programs from the pulse program library (TopSpin 1.3pl10) at 298 K. Chemical shift perturbations of HSQC peaks, upon addition of E peptide, were calculated with the following equation:

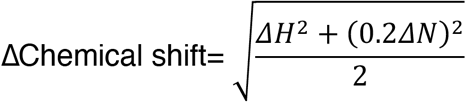

Dissociation constants (*K*_d_) were determined after the change in chemical shift was plotted against peptide concentration. Data were then fit to the following equation:

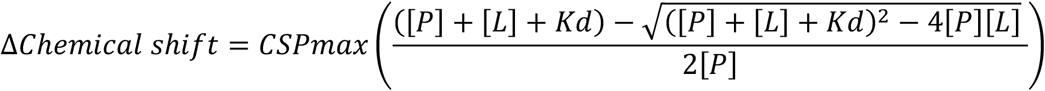

Where [P] is the concentration of BD2, [L] is the peptide concentration, *K*_d_ is the dissociation constant and CSPmax is the maximum chemical shift perturbation.

### Compound Treatments

Compounds, JQ1 (Selleckchem, S7110), dBET6 (Selleckchem, S8762), and ABBV-744 (Selleckchem, S8732), were dissolved in DMSO as per manufacturer’s instructions. Cells were treated at the time of infection for 48 hours with media changes with fresh compound-containing media every 24 hours.

### Compound Cytotoxicity Measurements

A549-ACE2 and Calu3 cells were seeded into 96-well plates and treated with identical compound concentrations used in the infection assays for 48 hours, where fresh compound-containing media was added every 24 hours. At the end of 48 hours, cell viability was assayed following the manufacturer’s protocol of CellTiter-Glo (Promega, G7571). Luminescence was recorded with an Infinite M Plex plate reader (Tecan) using an integration time of 1 second.

### Viral Infection Studies

A549-ACE2 and Calu3 cells were seeded into 12-well plates and rested for at least 24 hours prior to infection. At the time of infection, media containing compound and/or viral inoculum (MOI 0.01 or 0.1) was added on the cells for 24 hours. At 24 hours post infection, fresh compound-containing media or media only was added on the cells. The supernatant and cells were harvested by adding STAT-60 for downstream quantification of genes.

### Plaque-Forming Assays

Viral inoculations were harvested from experiments and serially diluted in DMEM (Corning). At the time of infection, the media on Vero-E6 cells were replaced with viral inoculation for one hour. After the one-hour absorption period, 2.5% Avicel (Dupont, RC-591 was layered on the cells and incubated for 72 hours. Then, Avicel was removed and cells were fixed in 10% formalin for one hour, stained with crystal violet for 10 minutes, and washed multiple times with water. Plaques were counted and averaged from two technical replicates.

### Animal Models

All protocols concerning animal use were approved (AN169239-01A) by the Institutional Animal Care and Use committees at the University of California, San Francisco and Gladstone Institutes and conducted in strict accordance with the National Institutes of Health *Guide for the Care and Use of Laboratory Animal (Council, 2011)*. Studies were conducted with male and female K18-hACE2 C57BL/6J mice (strain B6.Cg-Tg(K18-ACE2)2Primn/J) (The Jackson Laboratory, 034860). Mice were housed in a temperature- and humidity-controlled pathogen-free facility with 12-hour light/dark cycle and ad libitum access to water and standard laboratory rodent chow.

### Preparation of Compounds for Animal Study

A stock solution of JQ1 (50mg/ml in DMSO) and ABBV-744 (50mg/ml in DMSO) was prepared. JQ1 was then diluted to a working concentration of 5mg/ml in an aqueous carrier ((2-Hydroxypropyl)-β-cyclodextrin [Sigma C0926]) using vigorous vortexing. Mice were injected at a dose of 50mg/kg given intraperitoneally once daily. ABBV-744 was diluted to a working concentration of 5mg/ml in an carrier ((V/V): 2% DMSO, 30% PEG-400, 68% Phosal-50PG) using vigorous vortexing. Mice were treated at a dose of 20mg/kg given orally once daily.

### SARS-CoV-2 K18-hACE2 mouse infection model

A total of 50 animals were used in the study. Forty five animals were anesthetized and intranasally infected with 5×104 PFU/mL (50μl) of SARS-CoV-2 WA1 strain, five animals were mock infected and used as a control. The infected animals were divided in three groups each of 15 animals. Each group were treated with either JQ1 (50mg/kg) or ABBV-744 (20mg/kg), while DMSO at a vehicle concentration was used as a control. The animals were dosed from day 0 to day 7 post infection. All animals were monitored for their temperature and weight loss on daily basis. At 2 and 4 d.p.i. five animals from each group were euthanized by cervical dislocation and lung and brain tissue homogenized for plaque assays. Rest of the animals were monitored for their survival after infection. Lungs and cecum tissues from virus and mock infected animals were further processed for histological observations.

## QUANTIFICATION AND STATISTICAL ANALYSIS

The number of experiments and replicates are indicated in individual figure legends. Data were processed and visualized using GraphPad Prism. All quantified data are represented mean ± SEM or SD, as indicated, and quantification details are available in figure legends. Western blot band intensities were quantified using ImageJ.

## Acknowledgements

We thank all members of the Ott and Fujimori laboratories for sharing reagents, expertise, and feedback in the preparation of this manuscript. We thank Lauren Weiser and Veronica Fonseca for administrative support, Dr. Francoise Chanut for editorial support, and John Carroll for graphical support. This research is funded by grants from the National Institutes of Health: NCI R01CA250459 and NIAID R01AI137270 to D.G.F.; NIAID R37AI083139 to M.O. I.P.C. was supported by NIH/NIAID (F31 AI164671-01). S.M. was supported by TRDRP (T30DT1006) and the UCSF Discovery Fellowship. We are grateful for philanthropic support from QCRG philanthropic donors and the James B. Pendleton Charitable Trust. Imaging and image processing was performed at the Gladstone Institutes’ Histology and Light Microscopy Core. The SARS-CoV-2 (Isolate USA-WA1/2020) was deposited by the Centers for Disease Control and Prevention and obtained through BEI Resources, NIAID, NIH.

## Author Contributions

I.P.C., J.E.L., S.M., D.G.F., and M.O. conceived and designed the study. I.P.C., J.E.L. and R.K.S. performed experiments and data analysis. I.P.C. and R.K.S. designed and conducted the animal study. J.C.-S. and J.O. generated the knockout cell lines. M.Y.Z. contributed to binding studies. J.M.H., F.W.S., and T.Y.T provided experimental support. S.M. contributed to IP studies. M.G., M.Y.S., V.L.L., Y.L., Z.Y., E.W.T., A.D., and QCRG SBC provided protein expression support. D.G.F. and M.O. supervised the study design and provided technical guidance. K.H., N.K., D.G.F., and M.O. secured funding. I.P.C., J.E.L., D.G.F., and M.O. wrote the manuscript.

---

**Fig S1.**
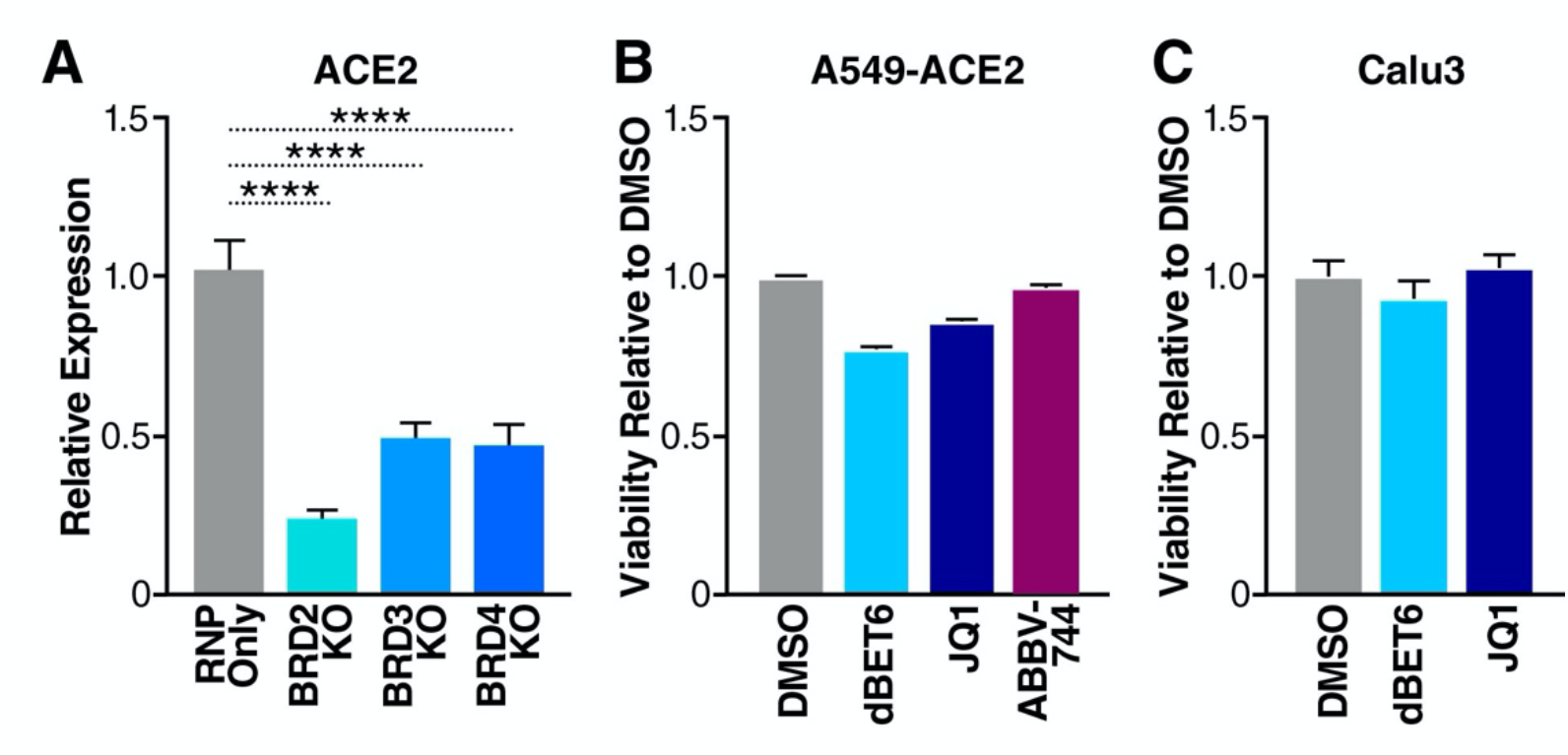
BET proteins are positive regulators of ACE2 expression. A. RT-qPCR of ACE2 RNA isolated from Calu3 cells with indicated BET KOs. Data are expressed relative to RNP Only cells. Average of three independent experiments analyzed in triplicate ± SEM are shown and compared to RNP Only samples by ANOVA: ****p<0.0001. B. Viability of A549-ACE2 cells treated with DMSO (vehicle), JQ1 (500nM), dBET6 (500nM), and ABBV-744 (500nM) for 48 hours relative to DMSO. C. Viability of Calu3 cells treated with DMSO (vehicle), JQ1 (500nM), and dBET6 (500nM) for 48 hours relative to DMSO.

**Fig S2.**
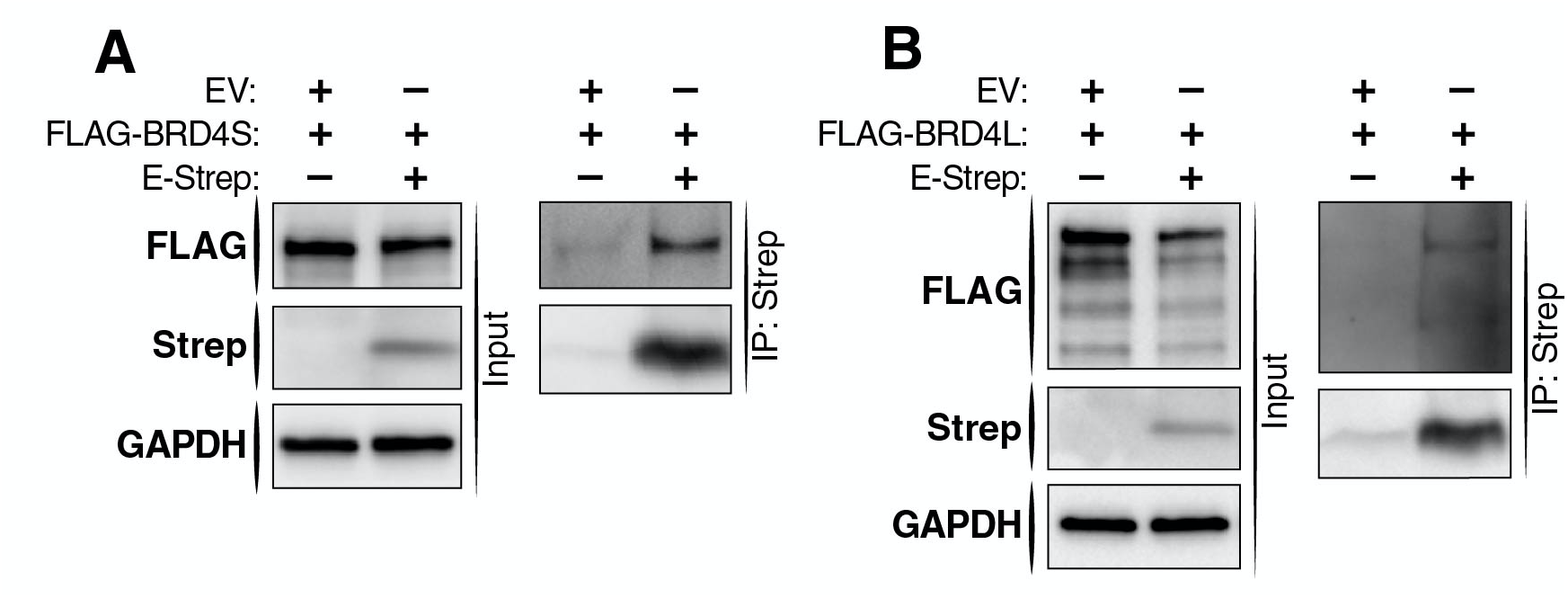
Both isoforms of BRD4 interacts with SARS-CoV-2 E protein. A. Immunoprecipitation of overexpressed Strep-tagged SARS-CoV-2 E protein (E-Strep) from transduced HEK293T whole-cell lysates, followed by western blotting using FLAG, Strep, and GAPDH antibodies. EV is empty vector control. B. Immunoprecipitation of overexpressed Strep-tagged SARS-CoV-2 E protein (E-Strep) from transduced 293T whole-cell lysates, followed by western blotting using FLAG, STREP, and tubulin antibodies. EV is empty vector control.

**Fig S3.**
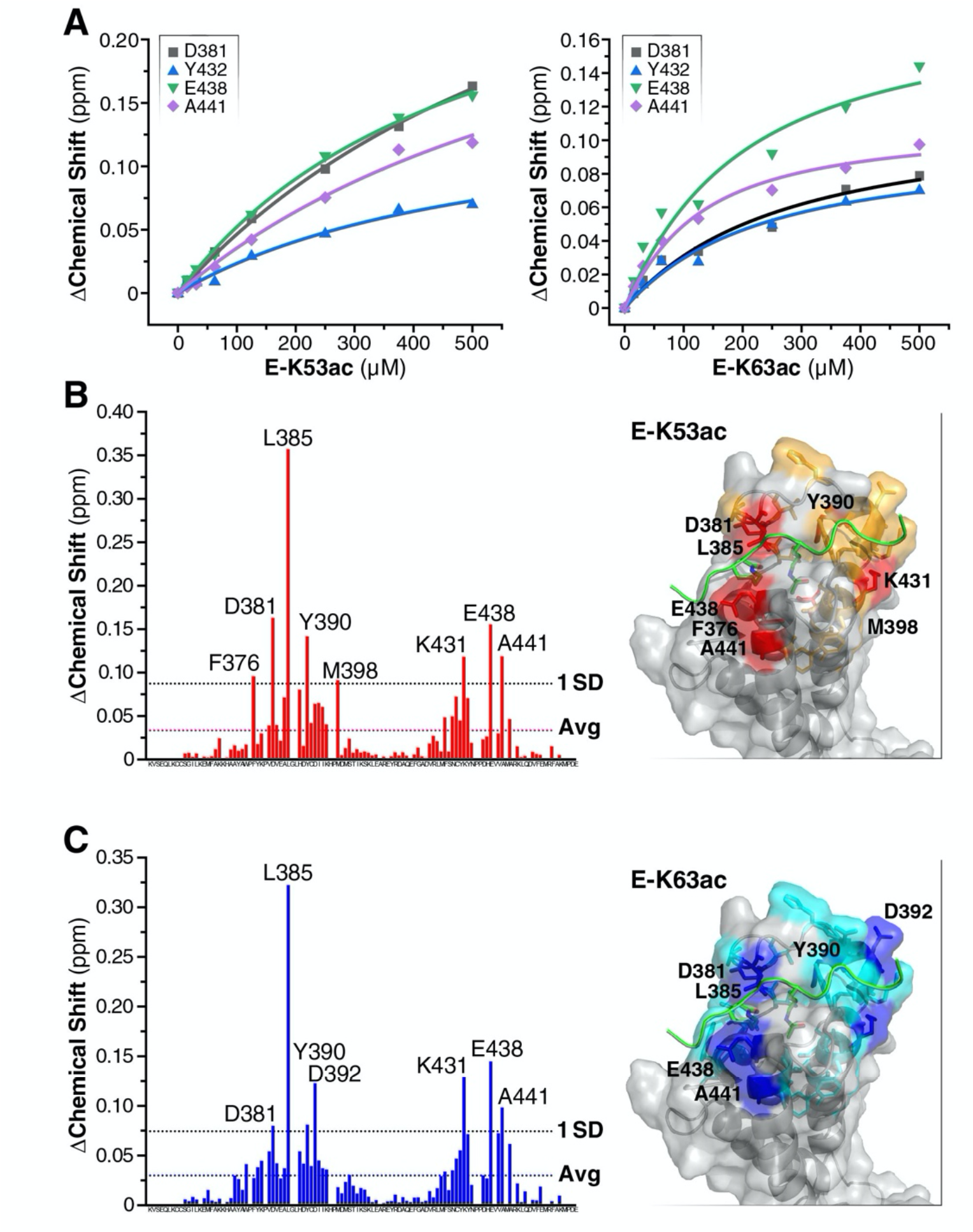
Acetylated E protein occupies similar binding sites to known acetylated lysine binding sites of BRD4 BD2. A. Binding curve of either E K53ac or E K63ac peptides binding to BD2 as measured by change in chemical shift. B. Change in chemical shifts in backbone amides of BD2 at a 1:10 molar ratio of protein to E K53ac (aa 48-58). The right-hand panel shows the structure of BRD4-BD2 in complex with Twist1 K73ac K76ac peptide (PDB ID: 2MJV). Twist1 is in green and acetylated lysine residues are shown. Residues shown in orange are those with a higher-than-average change in chemical shift and those shown in red have shifts larger than 1x SD. C. The same as B, but with E K63ac (aa 58-68) peptide. Residues shown in cyan are those with a higher-than-average change in chemical shift and those shown in blue have shifts larger than 1x SD.

**Figure S4.**
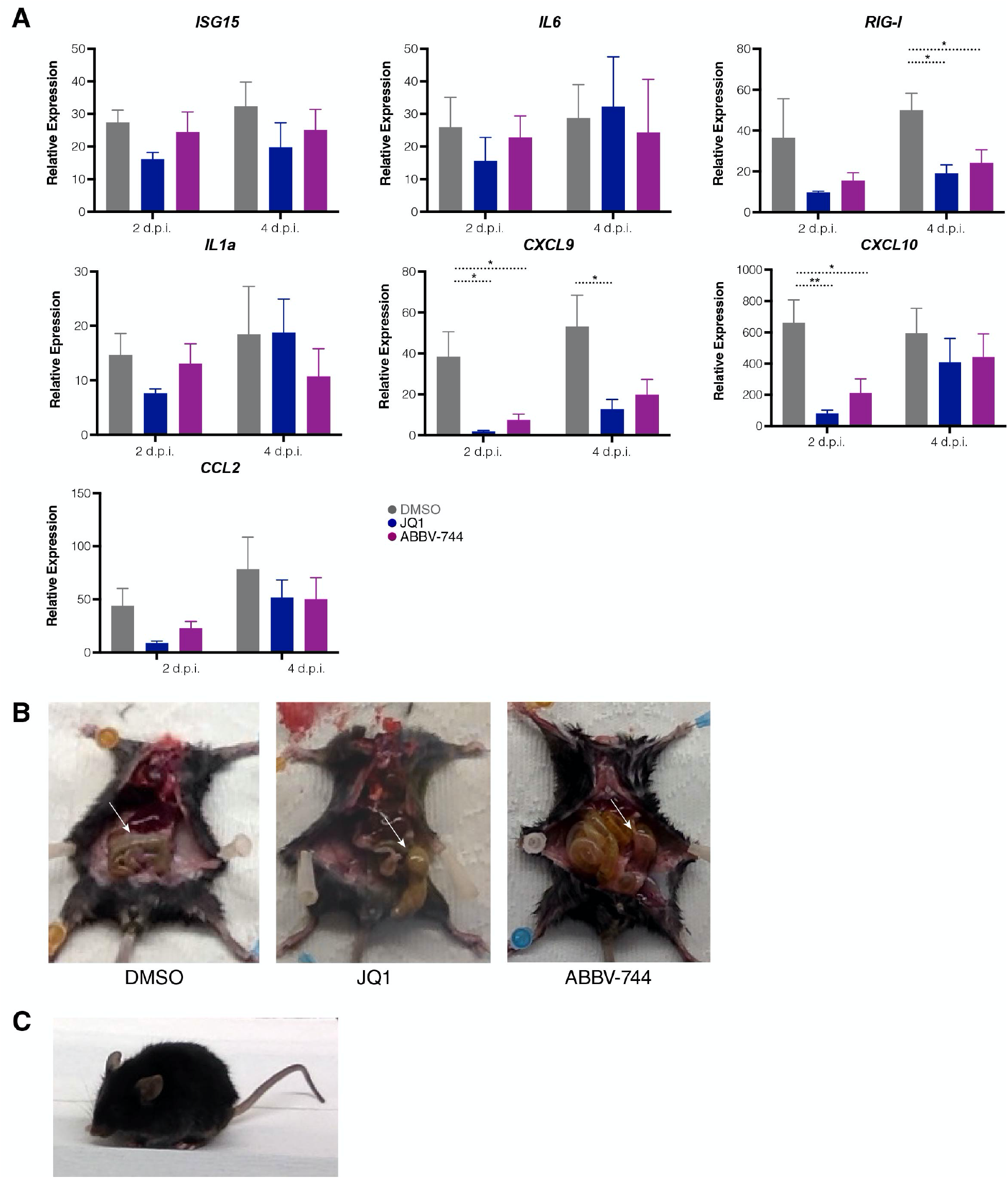
SARS-CoV-2 infection of k18-hACE2 mice results in dysregulated immune responses. A. RT-qPCR of RNA isolated 2 and 4 d.p.i. from lungs of SARS-CoV-2 infected mice treated with DMSO, JQ1, or ABBV-744. Data are expressed relative to uninfected mice for each respective gene. Average of 5 mice in each treatment group per time point analyzed ± SEM are shown and compared to DMSO by ANOVA: **p<0.005, *p<0.05. B. Representative images of the inflamed gut in SARS-CoV-2 infected mice. C. Representative image of an SARS-CoV-2 infected animal displaying the hunched back phenotype at 4 days post infection.

